# Visual neuroscience methods for marmosets: efficient receptive field mapping and head-free eye tracking

**DOI:** 10.1101/2020.10.30.361238

**Authors:** Patrick Jendritza, Frederike J. Klein, Gustavo Rohenkohl, Pascal Fries

## Abstract

The marmoset has emerged as a promising primate model system, in particular for visual neuroscience. Many common experimental paradigms rely on head fixation and an extended period of eye fixation during the presentation of salient visual stimuli. Both of these behavioral requirements can be challenging for marmosets. Here, we present two methodological developments, each addressing one of these difficulties. First, we show that it is possible to use a standard eye tracking system without head fixation to assess visual behavior in the marmoset. Eye tracking quality from head-free animals is sufficient to obtain precise psychometric functions from a visual acuity task. Secondly, we introduce a novel method for efficient receptive field mapping that does not rely on moving stimuli but uses fast flashing annuli and wedges. We present data recorded during head-fixation in areas V1 and V6 and show that receptive field locations are readily obtained within a short period of recording time. Thus, the methodological advancements presented in this work will contribute to establish the marmoset as a valuable model in neuroscience.

## Introduction

The common marmoset (*Callithrix jacchus*) has recently gained enormous popularity as an emerging model for neuroscience (Servick, 2018). Marmosets combine several advantages as a model animal: they are small in size, have fast reproduction cycles, and their rich behavioral repertoire makes them ideal to study a variety of complex and social behaviors (Koski and Burkart, 2015; Miller et al., 2016; Stevenson and Poole, 1976).

Recently, there have been successes in creating transgenic marmosets as disease models (Sasaki et al., 2009; Sato et al., 2020; Tomioka et al., 2017b, 2017a) and in the use of genetically encoded calcium indicators (Park et al., 2016). Due to the relatively high reproduction rates and the fact that marmosets often give birth to twins, it is likely that the marmoset will become a viable transgenic primate model (Kishi et al., 2014; Mitchell and Leopold, 2015; Shen, 2013).

The marmoset is of particular interest for the field of visual neuroscience (Mitchell and Leopold, 2015; Solomon and Rosa, 2014). Marmosets rely strongly on their sense of sight and therefore have a highly developed visual system. This is reflected by the fact that a large fraction of their neocortex is dedicated to visual processing (Rosa et al., 2009) and by the occurrence of brain networks for the processing of complex visual objects, e.g. faces (Hung et al., 2015). Due to the lissencephalic nature of the marmoset cortex (Heide et al., 2020), many visual areas are exposed on the surface of the brain, making them directly accessible for neuronal recording and imaging techniques. This enables the investigation of high-level brain areas that do not have clear homologues in the rodent and are difficult to reach in larger primates.

Previous neurophysiological studies have provided substantial groundwork on various aspects of the visual system of the marmoset (for a review see Solomon and Rosa, 2014). However, the majority of visual experiments in marmosets have been performed under anesthesia, thus making it impossible to identify the neuronal circuits underlying visual behavior. Therefore, it is crucial to develop methods for monitoring and manipulating neuronal activity in the awake animal.

Studies in the awake marmoset have predominantly been performed under head fixation. This has been used for various recording and stimulation approaches, like electrocorticographic (Hung et al., 2015), extracellular microelectrode (Johnston et al., 2018; Remington et al., 2012), and intracellular (Gao et al., 2016; Gao and Wang, 2019) neuronal recordings, microstimulation (Selvanayagam et al.,2019), fMRI (Belcher et al., 2013; Hung et al., 2015; Liu et al., 2013; Schaeffer et al., 2019), as well as calcium imaging (Mehta et al., 2019; Yamada et al., 2004) and optogenetics (Ebina et al., 2019; Macdougall et al., 2016). Psychophysical studies of marmoset vision have, to our knowledge, exclusively used head-fixed conditions, allowing precise eye tracking (Mitchell et al., 2015, 2014; Nummela et al., 2017). However, head-fixation can conflict with the execution of normal behaviors (Pandey et al., 2020; Populin, 2006), exposing the necessity for more naturalistic and less constrained paradigms (Krakauer et al., 2017; Sonkusare et al., 2019). Such paradigms have recently been successfully used to study the motor system (Kondo et al., 2018; Mundinano et al., 2018; Umeda et al., 2019), spatial navigation (Courellis et al., 2019), and the auditory system (Eliades and Wang, 2008; Roy and Wang, 2012) of the marmoset. Similar to the experimental setup, the design of stimuli can also be guided by the intrinsic behavior of an animal in order to gain understanding about brain function (Knöll et al., 2018).

The overall goal of this work was to adapt methods for visual neuroscience, such that they are more suitable for the behavioral requirements of the marmoset. We applied this idea to two key methods: Eye tracking and receptive field mapping. First, we remove the necessity for head fixation in visual psychophysics experiments and show that eyetracking quality from head-free animals is sufficient to measure precise psychometric functions. Second, we present a novel adaptation of visual stimuli for efficient mapping of receptive fields and provide neuronal data from areas V1 and V6 recorded under head-fixation.

## Results

### Eye tracking and behavioural setup

We used a standard infrared-based eye tracking system in combination with a custom calibration procedure to track the position of one eye of marmosets without head-fixation (Fig. 1a). Animals were trained to enter a tube-shaped chair and to position their head in front of a monitor without any immobilization of their body or head. The small size of the opening for the head allowed the animals to return to the same overall position and distance across behavioral sessions. Additionally, the reward-delivering lick spout was positioned centrally relative to the monitor, and animals quickly learned to keep their heads facing forward. This allowed successful tracking of pupil and corneal reflex as long as the animals were engaged in the task (Fig. 1b).

**Figure 1.**
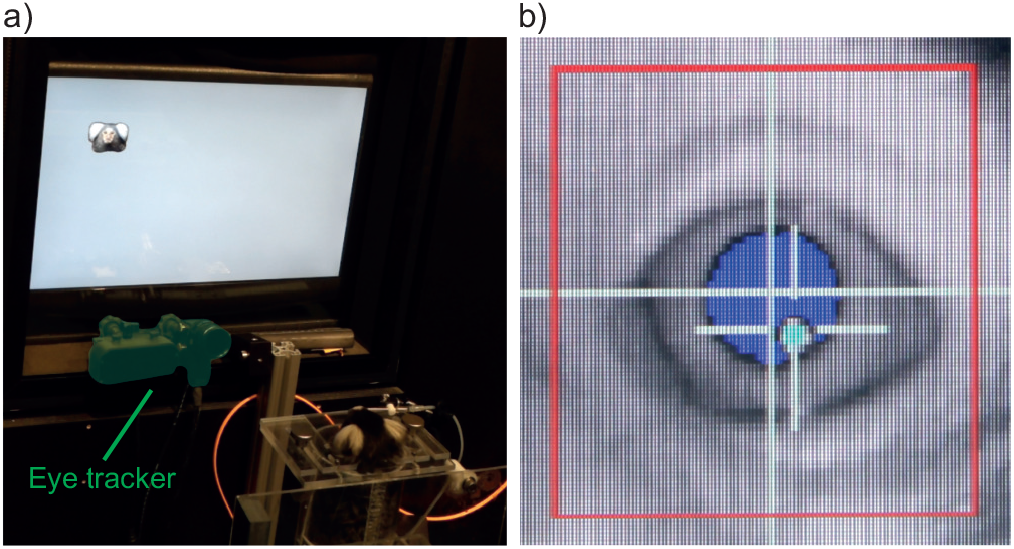
Head-free eye tracking setup. **a)** Experimental setup with a marmoset performing a simple face detection task. Eye tracking camera and IR-light source are highlighted in green. **b)** Online view of the eye tracking software. Pupil and corneal reflex are being tracked.

### Calibration

In studies with human subjects and nonhuman primates, calibration of the eye signal is typically performed at the beginning of each experimental session. This is done by a short sequence of fixations at defined coordinates on the monitor. This approach consumes time within each session, and provides only one or few data points per target position. Therefore, we introduced a different approach that uses average data from several trials performed during an entire calibration session. The resulting calibration was then used in future sessions, without any further corrections or offset removal. Figure 2a shows an example of such data after calibration. In a given calibration session, animals were required to briefly fixate a central fixation point and then saccade to a peripheral location indicated by a small stimulus (see Materials and Methods for details). The recorded eye data were analyzed offline together with the known target and fixation locations to create a calibration template for future sessions. Target locations with the highest density were manually selected and used to fit a third-order 2D polynomial function. The resulting transformation function was applied to the horizontal and vertical component of the eye data. The corrected positional data can be seen in Figures 2a and b. Figures 2c and d show calibrated example saccades towards different target locations. The density of eye data was particularly high at the central location (Fig. 2b). This is due to the fact that animals were required to initiate trials by maintaining their gaze at the central fixation point. This provided a large amount of data from various trials.

**Figure 2.**
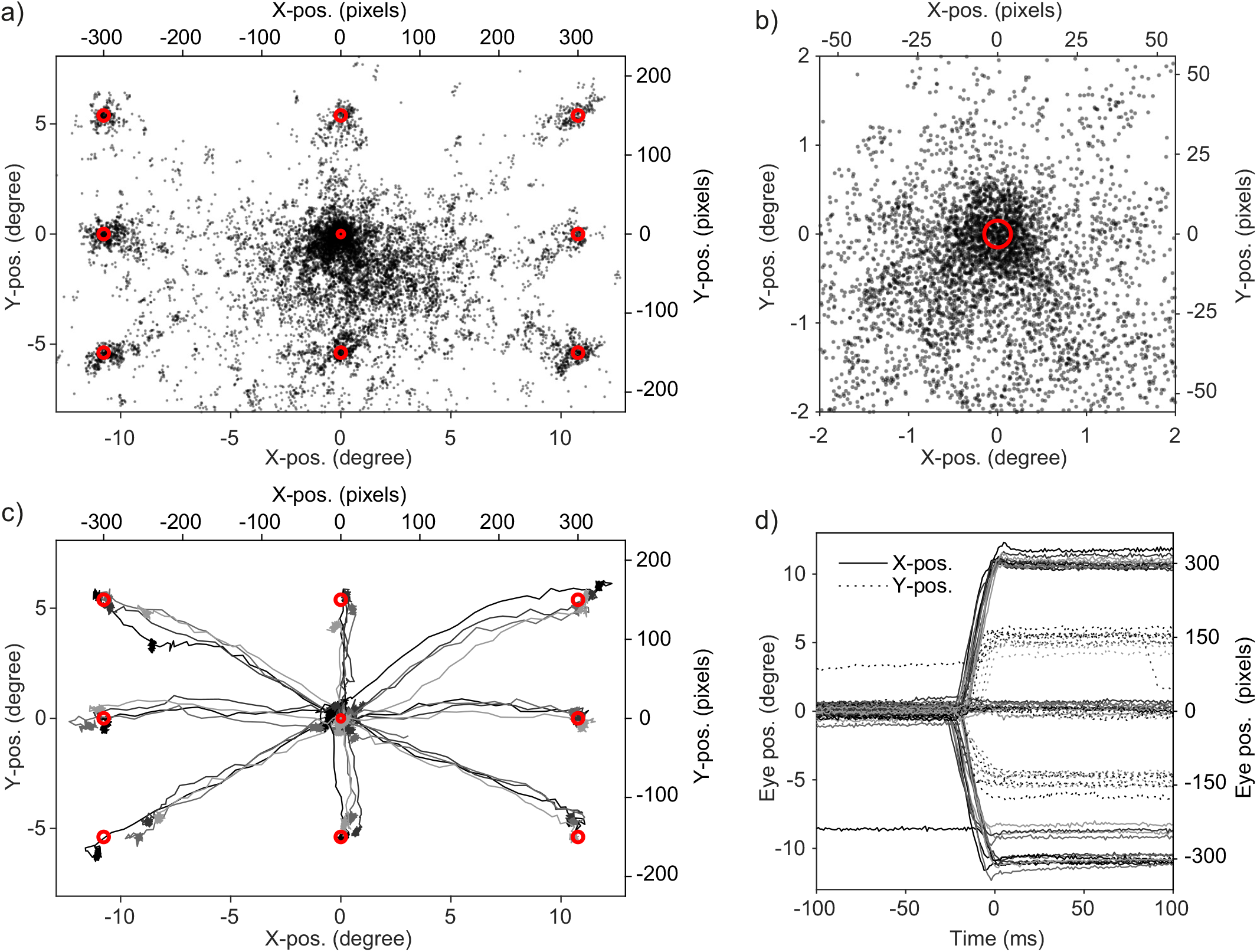
Offline calibration approach. **a)** Scatter plot of eye positions from an example calibration session. Every 25th sample is shown with 50% transparency for better visibility. Red circles indicate six times the standard deviation of the size of the visual stimuli used in the task. **b)** Same data as in a) but only showing the central 2 degrees. **c)** Saccade traces of four example trials per target position from the same data. **d)** Same data as in c) but plotted as a function of time for X-and Y-position separately. Data is aligned to the moment when gaze position entered the correct target window. Note that the example trials (first four trials per condition from an example session) include one trial with a double saccade and three trials with under-shooting saccades.

The obtained calibration was used in subsequent sessions to assess visual acuity, and several example trials from one such session are shown in Extended Data Movie 1.

### Accuracy and precision of central eye position

Offline calibration might be prone to inaccuracies resulting from changes in the position of the animal within and between sessions. We quantified this by calculating offset and standard deviation (referred to as ‘sigma’) for every session. We focused the analysis on the central 2.5 degrees and on the time from fixation onset until the animal made a correct saccade. This choice was motivated by the fact that the majority of studies use central fixation, and the presented procedure was primarily aimed to be used in such study designs. The density plots in Figures 3a-c show example sessions from each of the four animals. The binned and averaged horizontal and vertical eye data (Fig. 3a-c, black dashed lines) were fitted with 1D Gaussians (Fig. 3a-c, red lines), and those were subsequently used to derive horizontal and vertical offset and sigma values.

**Figure 3.**
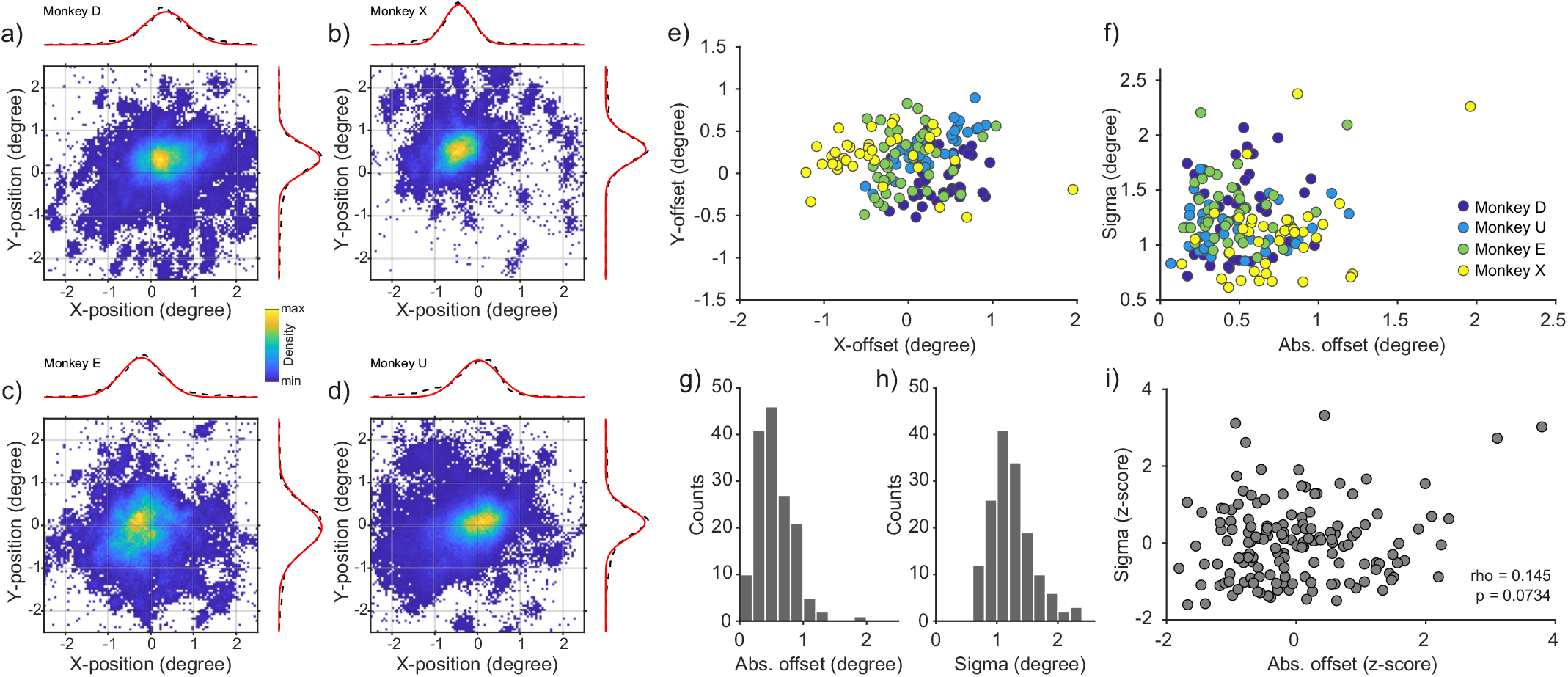
Quality of eye signals across sessions and animals. **a-d)** Example density plots of eye position during the visual acuity task for each monkey. Data are taken from time of fixation onset until correct response. Dashed lines show the average density for X-and Y-position. Red lines show Gaussian fits. **e)** X-and Y-offset values for all sessions (n = 153, color coded as indicated in panel f). **f)** Scatter plot of offset versus sigma values for all animals and sessions. **g), h)** Distributions of offset and sigma values for all sessions. **i)** Same as f), but after z-transformation per animal.

Figures 3e and f show the resulting offset and sigma values, respectively, for all 153 analyzed sessions. Overall, offset positions appear to spread around the fixation point. However, individual animals seem to form clusters (for example Monkey X in the top left direction), indicating small systematic biases at the level of individual animals (see Discussion). Most sessions showed an absolute offset that was below 1 degree (median = 0.503 ± 0.29 degree) and a sigma of less than 2 degrees (median = 1.17 ± 0.34 degree). As expected, these values were larger than measurements obtained during head-fixation (Extended Data Fig. 3-1 a-d; median of absolute offset = 0.177 ± 0.068 degree; median of absolute sigma = 0.430 ±-0.035 degree; Monkey D: n = 9 sessions; Monkey A: n = 8 sessions).

It is possible that the sessions with the largest offset would have the largest standard deviation. This might be expected in sessions in which the position of the animal is not optimal. To quantify this, we calculated the correlation between absolute offset and sigma values. To avoid any spurious effects of potential across-subject correlations, the offset and sigma values were first z-transformed per animal, and then combined before the correlation analysis (Fig. 3i). There was no significant correlation (Pearson’s rho = 0.145, p = 0.0734^a^), indicating that sessions with a large offset could still result in a reliable but shifted estimate of the eye position around the fixation point.

### Visual acuity in head-free marmosets

In order to test whether the head-free eye tracking quality is sufficient for visual neuroscience applications, we let animals perform a visual acuity task (Fig. 4). Animals could initiate trials by moving their gaze to a central fixation point. After 350-800 ms of fixation, a small Gabor stimulus (50% contrast) with variable spatial frequency appeared randomly at one of eight equi-eccentric locations. Trials were categorized as hits if the animal made a saccade to the target stimulus within 500 ms.

**Figure 4.**
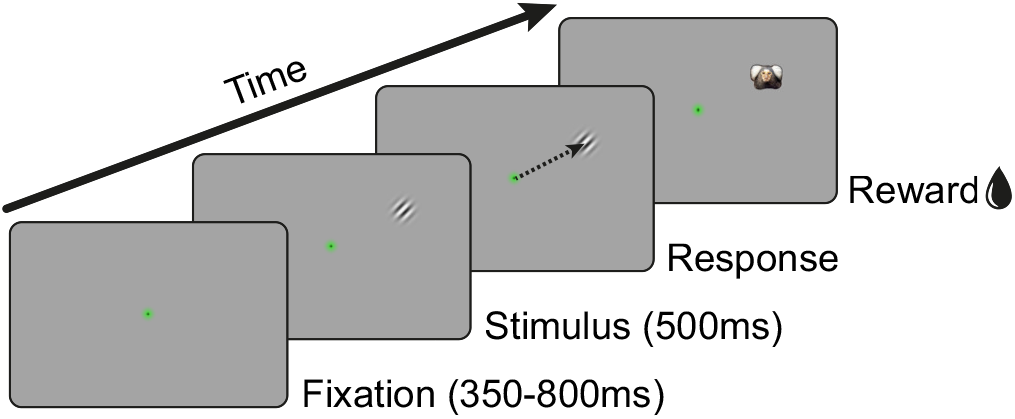
Visual acuity task design. Animals were required to fixate a central fixation point for 350-800 ms. A small Gabor stimulus (0.3 degree, 50% contrast) with variable spatial frequency was presented for 500 ms randomly at one of eight possible locations (*≈*10.77 degrees eccentricity). After a correct saccade to the target, a picture of a marmoset face was displayed and the animal was rewarded.

At the time of visual acuity measurements, all animals were familiar with simple face-detection tasks (Fig. 1a) and calibration tasks (Fig. 2) and had no problem switching to the visual acuity task. Animals performed on average approximately 100 hits per session. We fitted reaction times (RTs) and hit rates, as functions of the spatial frequency, with a four-parameter logistic function, separately per animal (Fig. 5a). Figure 5b shows the underlying RT distributions from hit trials across all conditions and sessions. RT values were longer for higher spatial frequencies: the average upper asymptote (245.5 ± 6.6 ms, mean ± SEM) was significantly above the lower asymptote (132.4 ± 2.7 ms; p = 0.00028^b^, paired t-test across the four animals). For the fits of the hit rates, the asymptotic value for high spatial frequencies was fixed to the chance level of 12.5%. The parameter that determines the inflection point of the psychometric function corresponds to the perceptual threshold. We calculated thresholds from hit rates and from RTs for each animal. As expected, thresholds were higher, corresponding to a higher visual acuity, when calculated from hit rates (6.4 ± 0.3 cycles/degree, mean ± SEM) as compared to RTs (5.2 ± 0.4 cycles/degree, mean ± SEM), indicating a more accurate estimation (Fig. 5c; p = 0.012^c^, paired t-test across the four animals). The thresholds obtained from hit rates are in line with previously reported values from Nummela et al. (2017) under head fixation. This confirms that it is possible to measure visual acuity in marmosets without head-fixation.

**Figure 5.**
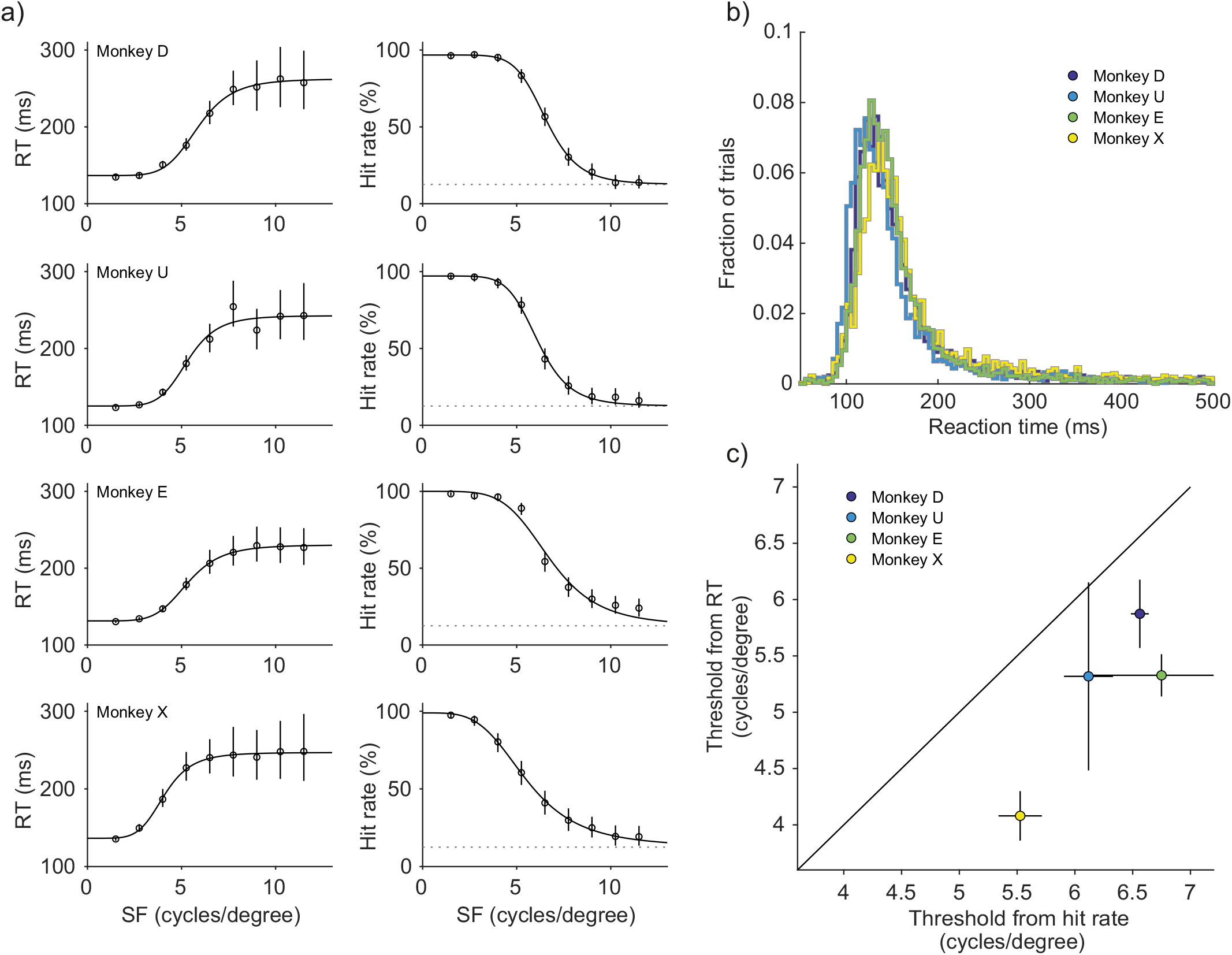
Visual acuity measurements in head-free marmosets. **a)** Reaction time (RT) and hit rate as a function of stimulus spatial frequency (SF) for each of the four animals. Mean RTs are shown on the left panel and hit rates are shown on the right panel. Error bars indicate 99.9% confidence intervals. In the hit rate plots, dotted lines show expected chance level performance of 12.5%. **b)** RT distributions, per animal, pooled over all conditions and sessions. **c)** Comparison of acuity thresholds calculated from RTs and hit rates. Error bars indicate 95% confidence intervals.

### Efficient receptive field mapping

Having established that marmosets can be behaviorally assessed without head fixation, we turned our interest to another important aspect of visual neuroscience, namely the mapping of visual receptive fields (RFs). As was the case for our head-free psychophysics experiments, the goal was to adapt a technique for a fundamental aspect of visual neuroscience in order to make it more suitable for the marmoset. Due to the small size of RFs in early visual areas, animals were head-fixed for the recordings (however, see Discussion). Commonly used stimulus types for RF mapping are: spatial noise (Citron and Emerson, 1983; Niell and Stryker, 2008), flashing dots or squares (Jones et al., 1987; Martinez et al., 2005; Tolias et al., 2001) and moving bars (Fiorani et al., 2014; Hubel and Wiesel, 1962). Another possibility is the use of rotating wedges in combination with expanding and contracting annuli. The latter is often used in fMRI studies (Benson et al., 2018; Sereno et al., 1995). Moving bars and rotating wedges are efficient stimuli due to their spatial correlation. However, because they are very salient, it can be challenging for animals to maintain their gaze at the fixation point. We developed a novel approach that uses fast flashing annuli and wedges to cover a large part of the visual field, thereby being very efficient. Importantly, the presented stimuli are flashed for a short duration at a random location and are either symmetric to or connected to the fixation point. We reasoned that this design might make it easier for the marmosets to maintain fixation.

### Annulus-and-wedge RF mapping task

Stimuli were black wedges and annuli, presented on a gray background. Wedges subtended 9 degrees of polar angle and were presented in steps of 4.5 degrees of polar angle. Annuli had a width that corresponded to the width of the wedge at the eccentricity of the midpoint of the respective annulus (Fig. 6). As soon as the animal moved its gaze to the central fixation point, a sequence of nine to ten stimuli was flashed within a typical trial. Each stimulus presentation lasted for eight frames at a monitor refresh rate of 120 Hz, resulting in a presentation time of ≈67 ms. At the end of a correct trial, animals were rewarded, and a marmoset face indicated the end of the trial. In case the animal broke fixation before the full stimulus sequence was over, we continued presenting the stimulus sequence and no reward was given. This was done to let the animal freely explore the stimuli in order to learn that there was no reward associated with looking at them.

**Figure 6.**
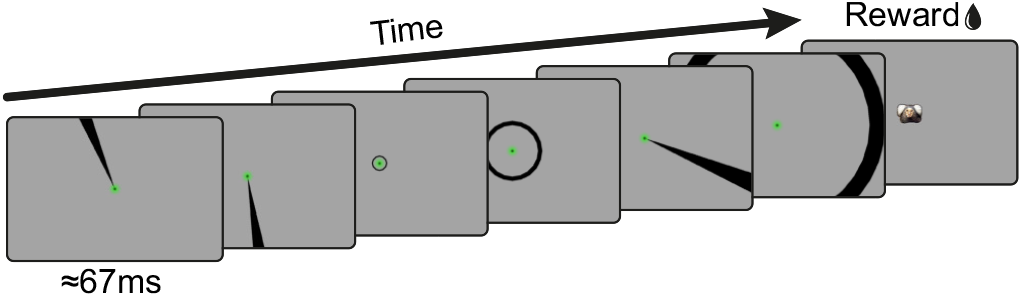
Receptive field mapping task. Annulus and wedge stimuli with sizes and orientations that covered the whole monitor were flashed for 8 frames (*≈*67 ms) each. The animal was required to maintain fixation throughout the trial and was rewarded at the end. Correct trials were signaled by the appearance of a marmoset face at the center. Typically, nine to ten stimuli were flashed per trial, but only six are shown here for clarity.

In order to test whether the RF mapping stimuli could be used to efficiently locate the positions at which neuronal activity was highest (i.e. location of the classic RF center), we recorded spiking activity from areas V1 and V6 (Fig. 7a). Animals had no problem maintaining fixation throughout many trials, and successfully completed several hundred stimulus presentations in short recording sessions (data from Fig. 7: n = 1654 in ≈11 min for Monkey D and n = 2529 in ≈12 min for Monkey A; corresponding to 9 to 17 and 14 to 24 repetitions per stimulus condition, respectively, after artifact rejection). We observed spontaneous neuronal activity on several recording sites as well as clear stimulus-evoked activity (V1 example trace in Fig. 7a). In total, 188 out of 384 (*≈* 49%) sites were significantly modulated from baseline and selected for further analysis (Monkey A: 63 of 64 sites in V1 and 60 of 128 sites in V6; Monkey D: 58 of 64 sites in V1 and 7 of 128 sites in V6).

**Figure 7.**
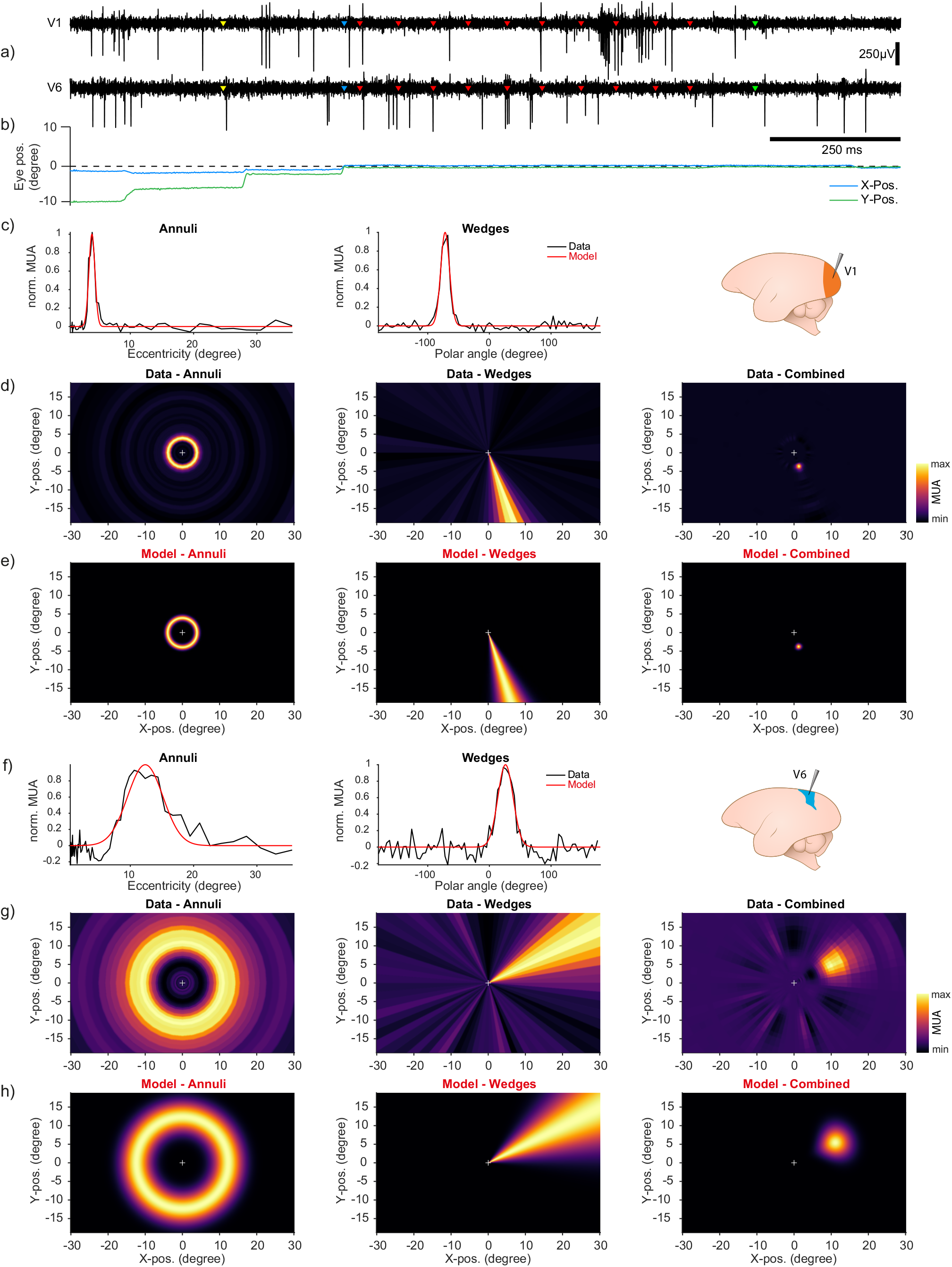
Receptive field mapping results. **a)** Spiking activity from two example channels simultaneously recorded from cortical areas V1 and V6 in Monkey A. Yellow markers indicate start of the trial. Blue markers indicate fixation onset. Red markers indicate onsets of individual stimuli. Green markers indicate time of reward. **b)** Corresponding eye trace for example trial. Note fixation onset as indicated by blue makers. **c)** Multi-unit activity (MUA) profiles and model fits for all presented annuli and wedges from an example recording site in area V1. **d)** Reverse correlation RF maps across the monitor, from annulus, wedge and combined MUA data. Color indicates normalized multi-unit response. White cross indicates position of fixation point at the center of the monitor. **e)** RF maps across the monitor, from annulus, wedge and combined model data. **f-h)** Same as c-e) but from recording site in area V6 in Monkey D.

We calculated the mean multi-unit activity (MUA) for all presented annuli and wedges separately, which resulted in activity profiles across all tested eccentricities and polar angles (Fig 7c, f, black lines). To estimate RF size and position, we fitted a Gaussian function or its circular approximation (von Mises function) to the annulus and wedge data, respectively (Fig 7c, f, red lines). The extracted RF sizes in V1 and V6 were largely consistent with previously published data from anesthetized animals (Chaplin et al., 2013; Rosa et al., 1997; Rosa and Schmid, 1995; Yu et al., 2020), although V1 RFs appeared slightly larger (Extended Data Fig7-1 a, see also Discussion). Based on the electrode target coordinates, the RF locations from V1 were expected to be in the lower visual field, relatively close to the vertical meridian (Chaplin et al., 2013). V1 RFs from both monkeys were consistent with this (Extended Data Fig. 7-1 b, d), suggesting placement in area V1. In V6, RFs spread across the upper and lower visual field, consistent with the compressed retinotopic representation of this area (Yu et al., 2020). Furthermore, as expected from the retinotopic organization of both areas, RFs from nearby electrode sites showed similar RF locations (Extended Data Fig. 7-1 b-e).

In order to relate spiking activity to the visual space across the monitor, we applied a reverse correlation technique (Jones et al., 1987; Ringach, 2004), that resulted in two activity maps (see Materials and Methods for details). Each map shows the location of those annuli or wedges, respectively, that evoked the highest spiking activity in the respective recording site (Fig. 7d, g). The two maps were then combined by multiplication to reveal the peak RF location of the recorded neurons. Maps from the model fits were obtained by transforming the model data from polar coordinates to Cartesian coordinates for each pixel of the monitor (Fig. 7e, f).

To quantify the efficiency of our RF mapping technique, we calculated the signal-to-noise ratio (SNR) for RF maps from a subset of the data. Subsampling was performed by randomly picking a small number of stimulus presentations per condition from the full dataset. This procedure was performed 100 times for each step, from one to nine repetitions. SNR was calculated as the amplitude ratio in decibels (dB) of the mean values inside and outside a region of interest (ROI) on the RF map. The ROI was determined by thresholding the RF maps that included all available data at the half maximum (see Materials and Methods for details). Figure 8a shows example RF maps calculated from a random small subset of stimulus presentations. The RF center position is already visible with one or two repetitions per condition and becomes clearer with more repetitions. As a baseline control condition, we also calculated RF maps from the stimulus-unrelated activity preceding the stimulus. The resulting RF maps show no spatially specific activation, due to the MUA being uncorrelated with the future stimulus. Accordingly, the SNR for the baseline condition remains flat, even when the number of repetitions increases (Fig 8b). In contrast, the SNR calculated from stimulus-evoked MUA increases with the number of repetitions. Crucially, SNR values were already high when a small number of repetitions was used.

**Figure 8.**
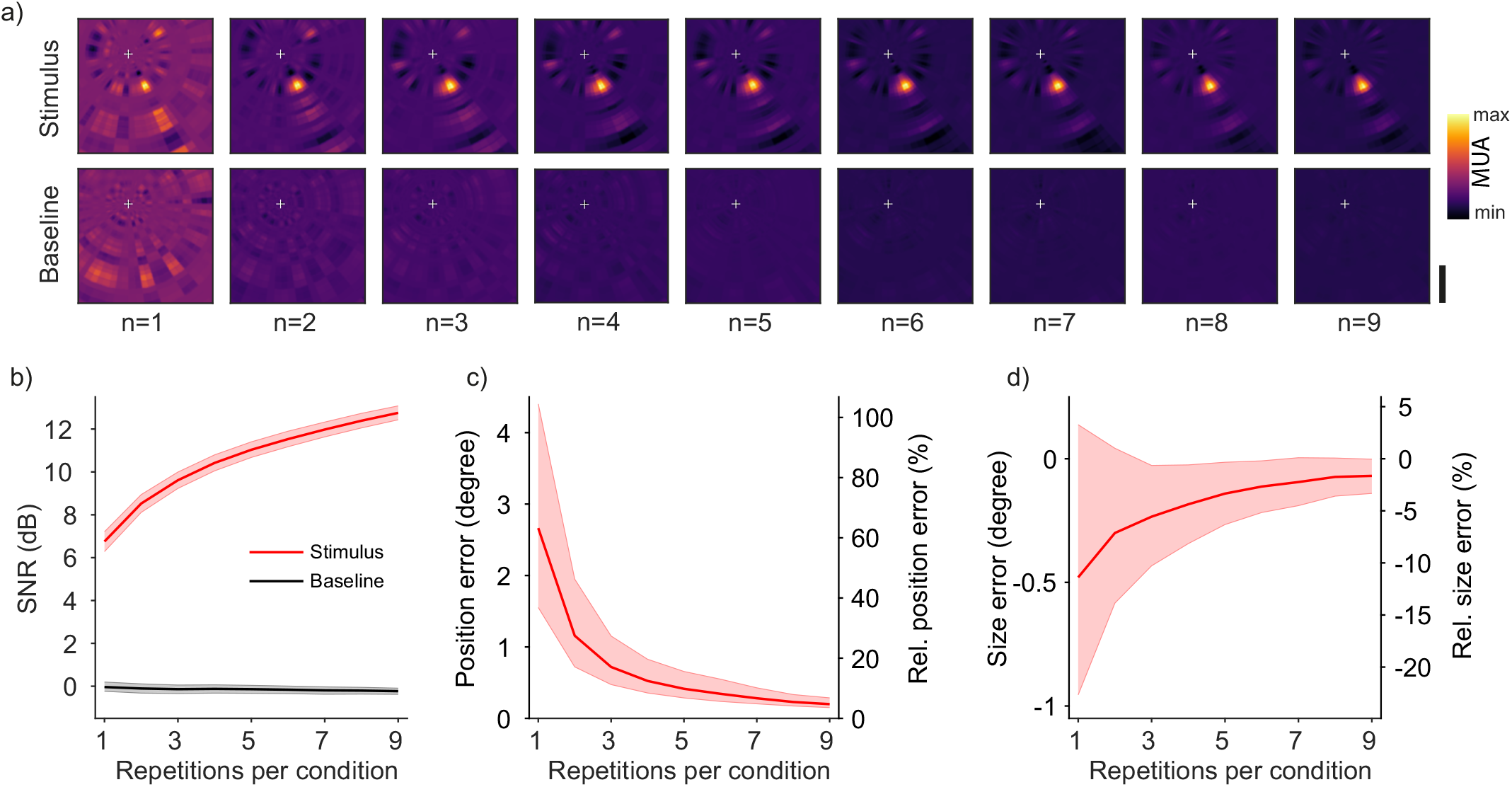
Efficiency of receptive field mapping. **a)** Example receptive field (RF) mapping results from subsampling procedure for up to nine repetitions per stimulus condition. Data from example recording site in Monkey D. RF location is already visible with a very low number of repetitions. White cross indicates position of fixation point at the center of the monitor. Color indicates normalized multi-unit response. Scale bar = 5 degrees. **b)** Signal-to-noise ratio (SNR) from subsampling procedure of RF mapping data, averaged across recording sites from both monkeys (n = 188). Reverse correlation analysis was performed from neuronal data following the stimulus presentation (red line) and, as control, preceding the stimulus presentation (black line). **c)** Position error and **d)** size error from subsampling procedure of RF mapping data, averaged across all recording sites from both monkeys. Shaded regions indicate 99.9% bootstrap confidence intervals.

Next, we quantified the absolute and relative errors in the estimation of RF position and size. We used the same subsampling procedure as in the SNR analysis and repeatedly calculated position and size of the resulting RFs with varying amounts of data. We then compared the subsampled results to the values obtained by using all available data. Relative errors were calculated by normalizing the absolute error to the size of the corresponding RF obtained from all available data. The errors for position and size converged quickly towards zero with relatively few repetitions per condition (Fig. 8c, d). The negative values for size errors reflect an initial underestimation of the size for small repetition numbers.

These results confirm that our annulus-and-wedge RF mapping technique is a very efficient way of mapping visual receptive fields. It yields reliable results with less than 10 repetitions per stimulus condition, which can be obtained in 5-10 min of recording time.

## Discussion

The goal of this work was to develop two key methodological advancements for visual neuroscience that are designed to be more suitable for the marmoset monkey. First, we show that head-free eye tracking in marmosets can be achieved with existing hardware. The analysis of accuracy and precision revealed that head-free eye tracking is potentially suitable for a variety of visual neuroscience applications. We tested the applicability by measuring visual acuity in four head-free marmosets and demonstrate that the obtained acuity thresholds are consistent with previously published data obtained under head fixation (Nummela et al., 2017). Second, we introduce a novel method for efficient receptive field (RF) mapping that does not rely on moving stimuli, but uses a sequence of rapidly flashing annuli and wedges. To validate the novel RF mapping technique, we recorded data in areas V1 and V6 and show that RF locations are readily obtained in a short time and with very few stimulus repetitions.

### Limitations and advantages of head-free eye tracking and offline calibration

As expected, the accuracy and precision obtained from head-free animals is lower than what can be achieved under head fixation (Extended Data Fig. 3-1 a-d). Some of the imprecision can be explained by our choice of lens that was not optimized for maximum magnification of the pupil into the field of view (Fig. 1b). Instead, the lens was chosen to have a large field of view (i.e. a large ‘head box’) that allowed capturing pupil and corneal reflex signals despite head displacement. A large part of the inaccuracy arises most likely from movement of the animals within a session and across sessions. In principle, these offsets and errors from the resulting head-rotation could be partly compensated by tracking the animals’ head position in real time (Niehorster et al., 2018; SR Research, 2009). However, this can require the placement of a physical tracking target on the subject (Wang et al., 2019), which might be difficult to apply when working with small animals. More complex, model-based algorithms for head tracking (Yang and Zhang, 2002) might resolve this, and commercial systems (albeit optimized for humans) do already exist (Niehorster et al., 2018). The approach presented here focused on the implementation of a procedure with minimal hardware and software changes. It can therefore be used within the configuration range of existing commercial eye trackers in many laboratories.

Inaccuracies in eye position measurements might influence the outcome of a study (Hessels et al., 2016; Holmqvist et al., 2012). It was therefore crucial to validate our approach against previously reported data. The task described here was used because it had been successfully deployed before in head-fixed marmosets to assess visual acuity (Nummela et al., 2017). This allows for a close comparison of results and confirms that it was possible to measure visual acuity thresholds without the need for head fixation. It should be noted however that small differences in the experimental setup (i.e. monitor specifications, luminance, and distance to the screen) could still result in different acuity thresholds. We also found that acuity thresholds were higher, corresponding to a higher visual acuity, when calculated from hit rates (Fig. 5c). This confirms that hit rates are the preferred measurement for the estimation of perceptual thresholds (Palmer et al., 2005).

As mentioned above, offset values from individual animals appeared to cluster around a non-zero value across sessions (Fig. 3e), e.g. sessions from Monkey X seemed to cluster in the top left direction. Such clustering potentially indicates small biases during the calibration session. If so, offsets might be further reduced by subtracting the average offset across all recorded sessions, and thereby posthoc recalibrating the data. This could also be used to implement an iterative procedure that uses every recorded session to optimally compensate for offsets.

There are however limitations to the precision that can be achieved with the head-free eye tracking approach presented here. Most sessions showed an absolute offset (analyzed within the central 2.5 degree) that was below 1 degree (median = 0.503 ± 0.29 degree) and a standard deviation (sigma) of less than 2 degree (median = 1.17 ± 0.34 degree). These values indicate that it would not be feasible to perform experiments requiring continuous high-precision eye-position control. Nevertheless, the data quality is sufficient for many psychophysics applications (see below) and might even be used in experiments in which neuronal recordings are made from brain areas that have large receptive fields and exhibit translation invariance, such as the inferior temporal cortex (Rolls et al., 2003; Tovee et al., 1994).

We used nine calibration points and a polynomial function to calibrate eye data. The calibration function relies on interpolation between those points and extrapolation beyond the points. Therefore, care should be taken about the choice of the calibration points and the interpretation of eye data beyond the calibration points. Our procedure made use of the relatively long fixation times at the central fixation (Fig. 2b). This sampled eye data across varying head positions, and thereby resulted in a robust estimate for the calibration around the central location. In the same way, one could optimize the calibration positions to match the future positions for any task, thereby minimizing the effects of interpolation errors. Practically, this would entail that the very stimulus positions used in the actual task are also used for calibration. Furthermore, increasing the number of calibration targets or using smooth pursuit eye data can improve accuracy (Hassoumi et al., 2019; Kasprowski et al., 2014; Pfeuffer et al., 2013).

Head movements during the task will influence eye-tracking quality and can result in loss of signal (Niehorster et al., 2018). The eye tracker used in this study allows for a maximum of ±25 mm horizontal or vertical head movement without accuracy reduction (SR Research, 2009). The horizontal limits and the upper vertical limit are most likely not reached because the small opening for the head does not allow such large movements. From our observations, we estimate the actual possible head movement to be approximately ±15 mm. The lower vertical limit might be reached when the animal is retracting its head partly into the chair. Other movement types (i.e. yaw, roll and pitch) will also contribute to the reduction of signal quality and have been described in detail for human subjects (Ehinger et al., 2019). Furthermore, head-movement related changes in the distance between the eye and the camera can result in erroneous changes in calibration gain. An increased gain will make it more difficult for the animal to maintain fixation. It might also result in “overshooting” when executing a saccade to the target position. A gain-increase can therefore result in more aborted trials (“break fixation” trials) and in fewer correct trials due to “overshooting”. In the case of a gain decrease, maintaining fixation might be easier for the animal, due to the apparently lower amplitude of eye movements. However, correct execution of a saccade to the target position might be impaired due to “undershooting” arising from the lower gain. In general, “undershooting” or “overshooting” are not likely to cause large changes in trial outcome because the target tracking windows used in this study are relatively large (3-4 degrees radius). Importantly, such changes would also affect the easiest conditions, thereby decreasing the upper asymptote of the psychometric curves. However, all animals in this study have a hit rate close to 100% for the easiest stimuli (Fig. 5a). This is an indication that there was no strong influence resulting from changes in gain that could arise from the positional changes of the animals.

Changes in the distance between the eyes and the monitor, i.e. depth movement, would also lead to a change in stimulus position and size on the retina of the animal. An increased distance would result in lower eccentricity and could thereby increase the animals’ detection performance. At the same time, it would result in decreased angular stimulus size, thereby counteracting the performance increase to some extent. Since we did not measure head-position data, we cannot quantify how this might have influenced our results. However, we can provide an estimate on the expected maximum change in stimulus eccentricity and size, based on the geometry of our setup. Due to the small opening for the head and the position of the lick-spout in front of the animal, we estimate the actual depth movement to be smaller than ±5 mm. Under the very conservative assumption of ±10 mm depth movement, the calculated difference in stimulus eccentricity is ±0.23 degrees. The resulting change in angular stimulus size is ±0.024 degrees, in the opposite direction. Considering that stimuli are presented in the visual periphery (at 10.77 degrees), where eccentricity-dependent changes in visual acuity are not as steep as in the fovea, the influence of depth movements on our results are expected to be very small.

We focused our analysis mainly on the eye data around the central fixation location. This was primarily done to reduce the influence of variance arising from the behavior of the animals. Although we did not measure this explicitly, it can be expected that head movements are being executed when the animals need to position their gaze on a peripheral target (Pandey et al., 2020). Without head movement, marmoset eye position is most of the time within the central 5-10 degrees (Mitchell et al., 2014). Therefore, peripheral eye data, which is not precisely corrected for head position, should be interpreted with care. Yet, a large proportion of visual psychophysics studies do not require high accuracy eye-tracking in the visual periphery, but only in the center (Anton-Erxleben and Carrasco, 2013; Carrasco, 2011). The visual acuity task presented here is an example of such a task design. Target locations are far enough apart and their respective target windows can be relatively large, therefore allowing the animal to perform the task even under noisy conditions (see also Extended Data Movie 1). However, it is still important to track eye position reliably during the fixation period in order to prevent the animal from moving its gaze closer to the target. Such a change in eye position would bring the stimulus closer to the fovea, which in turn would result in a higher visual acuity (Chaplin et al., 2013; Nummela et al., 2017). This potential confound is additionally controlled for by using a randomized and symmetrical stimulus arrangement (Fig. 4).

The use of head-free eye tracking can provide several advantages. It enables the investigation of behaviors that are difficult or impossible to be studied under head fixation. Sound localization for example is strongly impaired during head-fixation (Populin, 2006) and the natural pattern of eye movements can be disrupted in some animals (Meyer et al., 2020; Wallace et al., 2013). Other behavioral tasks per definition require an unconstrained animal and can at best be approximated in virtual environments, i.e. tasks related to navigation, complex movement, social interactions and foraging. A further advantage of our approach is that it can be used in completely naïve animals, without the need for implantation of a head-post. This could be used for screening animals prior to implantation to select individuals with normal acuity (Graham and Judge, 1999) and overall good behavioral performance. Additionally, naïve animals can be pre-trained for complex tasks with an automatic home-cage training setup (Berger et al., 2018; Calapai et al., 2017), thus potentially increasing the number of animals in a study.

Under voluntary semi-automatic conditions, marmosets work reliably but for relatively short amounts of time (Walker et al., 2020), thus making every minute of recording time very valuable. We show that it is possible to perform psychophysical measurements without spending time on daily re-calibration. This reduces the stress for both the animal and the experimenter and increases data collection time. Future work might make use of fully automated experimental setups (Poddar et al., 2013), with multi-camera tracking (Young et al., 2016) and advanced 3D pose estimation (Mathis et al., 2018; Nath et al., 2019), potentially in combination with wireless neuronal recordings (Courellis et al., 2019; Eliades and Wang, 2008; Roy and Wang, 2012).

### Limitations and advantages of annulus-and-wedge RF mapping

RF mapping techniques are often optimized for specific requirements and scientific questions. Stimuli such as spatial noise patterns (Citron and Emerson, 1983; Niell and Stryker, 2008) and to some extent flashing dots or squares (Jones et al., 1987; Martinez et al., 2005; Tolias et al., 2001) can be used to infer detailed spatiotemporal RF characteristics (Ringach, 2004). However, due to the large number of possible stimulus configurations, such approaches can be time-consuming when RF centers need to be localized across large parts of the visual field. Moving bars (Fiorani et al., 2014; Hubel and Wiesel, 1962) and moving annulus-and-wedge stimuli (Benson et al., 2018; Sereno et al., 1995) are more suitable for that purpose due to their spatial structure. The stimulus design presented here was motivated by two factors: The behavioral requirements of the marmoset and the experimental requirement to locate RF centers with unknown positions and sizes across two visual areas. We initially presented some animals with moving bars and observed that they would break fixation and follow the bar movement on almost every trial, thereby making data collection nearly impossible. This comes as no surprise given that marmosets are prey animals and that it is essential for their survival in the wild to detect potential predators (Ferrari, 2008).

The flashing annulus-and-wedge RF-mapping approach combines several properties that are advantageous for the localization of RF centers with little data. The time for a single stimulus presentation is short (*≈* 67 ms), and the set of stimuli covers the entire monitor. This enables mapping of a relatively large part of the visual field within a short duration. The design of annuli and wedges corresponds to a polar coordinate system. Every position on the monitor will at some point display a wedge with a specific polar angle and an annulus with a specific eccentricity. In this way, for each neuron, or MUA, the polar angle and eccentricity for which it shows the maximum response can be determined (Fig. 7c-h). Yet, the stimulus set does not cover all orientations equally at every position. This might lead to a reduced response in case neurons are not optimally tuned to the annuli and wedges shown in their respective RFs. This limitation could be addressed by filling the stimuli with textures of randomized orientations.

One additional feature of the design is the stimulus scaling with eccentricity. The increasing RF size with eccentricity due to cortical magnification is a well-established phenomenon (Harvey and Dumoulin, 2011). This is one of the reasons why rotating wedges and contracting/expanding annuli are typically used in fMRI experiments where is it necessary to stimulate large visual cortical regions across eccentricities (Benson et al., 2018; Sereno et al., 1995). A related approach has been used by Hung et al. (2015) to map the foveal bias of visual areas with fMRI in the marmoset, albeit without strict eye fixation and the lack of appropriate stimulus resolution. In contrast, bar stimuli with a fixed width will be suboptimal both in driving neurons that have small parafoveal RFs and neurons that have large peripheral RFs. An RF-mapping stimulus that takes eccentricity into account will elicit stronger and more evenly distributed neuronal responses across the cortex.

The reported errors for RF position and size estimates were found to be small even after few repetitions per stimulus condition (Fig. 8c, d). RF positions and sizes from both areas were largely consistent with previously published data (Extended Data Fig7-1 a-e). Notably, the RF size estimates from area V1 appear to be approximately 0.5-1 degree larger than previously reported in anesthetized marmosets (Rosa et al., 1997). This discrepancy is most likely due to the fact that our results are obtained from awake animals that execute small eye movements (micro-saccades) during the fixation period. Additionally, our results are obtained from MUA, in which multiple neurons are pooled together, thereby also leading to larger RFs. However, we cannot exclude the possibility that the relatively large annuli-and-wedge stimuli contribute to an over-estimation of RF size. The precise influence of stimulus features on the resulting RF properties will need further detailed studies. Importantly, the estimation of RF location should remain mostly unaffected by this.

### Conclusion

Our work contributes to the rapidly growing field of marmoset monkey research. The concepts of less constrained paradigms and the adaptation of stimuli to the ethological needs of a species might be transferred to other species and to other areas of research. Together, this will promote diversification of the animal model landscape (Hale, 2019; Hemberger et al., 2016; Keifer and Summers, 2016; Yartsev, 2017) and solidify the contribution of marmoset research.

## Materials and Methods

All animal experiments were approved by the responsible government office (Regierungspräsidium Darmstadt) in accordance with the German law for the protection of animals and the “European Union’s Directive 2010/63/EU”.

### Animals

Five male adult common marmosets (*Callithrix jacchus*) were included in this study. Data from one other animal was excluded because the number of collected trials in the visual acuity task was much lower than in the other animals (Excluded animal: n = 1637 hits, vs. Monkey D: n = 4877 hits, Monkey U: n = 3918 hits, Monkey E: n = 4838 hits, Monkey X: n = 2951 hits). Animals were typically housed in groups of two or three. The housing area was kept at a temperature of 23-28 °C and at a humidity level of 40-70%. The dark/light cycle was 12h/12h, switching at 6:00/18:00.

### Food schedule and reward

Animals were fed in their home cages with marmoset pellets, nuts, fresh fruits and vegetables. Animals were on a mild food schedule and had *ad libitum* access to water. Typically, food was removed from the home cage after 17:00, and animals went into training/recording sessions the following day between 10:00 and 14:00. No large changes in body weight in relation to the training were observed. The reward during the tasks was a viscous solution of gum arabic (gum arabic powder, Willy Benecke, Germany) applied through a syringe pump (AL-1000HP, WPI, USA) that was controlled by a custom Arduino-based circuit. The amount of reward was typically between 0.05-0.09 ml per trial, and manually adjusted according to the performance of the animal.

### Behavioral training

Naive animals were first slowly acclimatized to be transported inside a dedicated transport box from the housing area to the laboratory setup. In the setup, they were allowed to enter the primate chair through a tunnel that connected the transport box with the chair. The position of the chair remained fixed across sessions, and the opening for the head was minimized (36 mm wide and 40 mm long), which assured that the head remained within a small region across sessions. As soon as the animals felt comfortable to stick out their head from the chair, they were rewarded manually through a lick spout that was placed in front of the animal’s mouth. The lick spout position was adjusted per session with regard to horizontal distance to the animal’s mouth to enable easy licking, yet the lateral and vertical position remained fixed across sessions. As the animals developed a stereotyped licking routine, this further contributed to constant head positioning during eye tracking. After this initial training, the setup was configured to automatically reward the animal whenever an eye signal could be detected. This led to the animal being conditioned to face forward and to look directly at the monitor. At this stage, animals were ready to be presented with visual stimuli that could be used to perform the initial eye calibration. For this we showed small marmoset faces at defined positions on the monitor (Fig. 1a).

### Stimulus presentation

Stimulus presentation was controlled by the custom-developed ARCADE toolbox (https://github.com/esi-neuroscience/ARCADE), based on MATLAB (Mathworks, USA) and C++. Stimuli were displayed on a TFT monitor (SyncMaster 2233RZ, Samsung, South Korea) at a refresh rate of 120 Hz. The monitor was gamma corrected and placed at a distance of 45 cm in front of the animal. Animals performed the task in a dimly lit recording booth. A photodiode was placed in the top left corner of the monitor in order to determine exact stimulus-onset times.

### Eye tracking

The left eye of the animals was tracked at 1 kHz sampling rate with a commercial eye tracking system (Eyelink 1000, SR research, Canada). Corneal reflex and pupil were tracked under external illumination with infrared light. A 25 mm/F1.4 lens was used at a distance of 28 cm to the animal’s eye. This resulted in a relatively large field of view, allowing for eye tracking despite head movement.

### Calibration and analysis of eye data

An initial coarse calibration via the Eyelink software was performed before using the system to collect data for offline calibration. For this, we used large salient stimuli (e.g. faces) presented at the default Eyelink calibration points (nine-point calibration). We manually accepted the position of the gaze to the target location with a key press. This initial coarse calibration was then used for each animal to perform the actual calibration task with large tracking windows (3-4 degrees). During the calibration task, a fixation point was shown that consisted of two overlaid Gaussians (one displayed in the background and colored green with a size of 0.15 degree standard deviation, the other one black with a size of 0.05 degree standard deviation). Animals were required to fixate the fixation point for 150-300 ms, at which time a small black Gaussian stimulus (0.08 degree standard deviation) was presented at one of nine possible calibration positions: 0/0, −300/-150, −300/0, −300/150, 0/150, 300/150, 300/0, 300/-150, 0/-150 pixels, referenced from the center of the monitor (Fig.2a). After a correct saccade to the target (reaching a window of 3-4 degrees around the target within 50-800 ms after target onset) and 100 ms of fixation on the target, a picture of a marmoset face was displayed and the animal was rewarded.

The uncalibrated eye data was plotted as shown in Figure 2a, and the positions with highest density were manually selected. The extracted coordinates from the selected eye positions and the known calibration points were then used to fit a third-order 2D polynomial function (“fitgeotrans” function in Matlab) to generate a template calibration for each animal.

For the estimation of accuracy (offset) and precision (sigma), 153 sessions from four marmosets were analyzed (Monkey D: n = 38 sessions, Monkey U: n = 37 sessions, Monkey E: n = 40 sessions, Monkey X: n = 38 sessions). Six sessions were excluded, because eye data was lost due to storage issues. For every session, eye data was binned with a bin size of 0.05 degree. The x and y components of the central 2.5 degrees were averaged and fitted with a 1D Gaussian function, separately per animal and session. The mean of the Gaussian fit corresponds to the x and y offset from zero. The Euclidean distance from zero to the X-and Y-offset-values from the fits was taken as the absolute offset. The Euclidean distance from zero to the x-and y-sigma-values from the fits was taken as the absolute sigma value.

### Visual acuity task

Animals were required to fixate a central fixation point for 350-800 ms. After this period, a small Gabor stimulus (50% contrast, 0.3 degree standard deviation in size, random orientation) was presented at one of eight possible equi-eccentric locations at a distance of 300 pixels (*≈* 10.77 degree) from the center. No corrective changes to the stimulus position were made. Spatial frequency values varied across trials and could assume the following values: 1.5, 2.75, 4.0, 5.25, 6.5, 7.75, 9.0, 10.25, 11.5 cycles/degree, with the probability of the lowest spatial frequency condition being twice as high as the other conditions. Trials were categorized as hits if the animal made a saccade to this stimulus within 500 ms. Responses that were faster than 50 ms were categorized as early responses and were not rewarded. After a correct saccade to the target, a picture of a marmoset face was displayed and the animal was rewarded. The amount of reward was typically between 0.05-0.09 ml per trial, and manually adjusted according to the performance of the animal. Animals were rewarded with a small reward (0.0025 ml) when they missed the target but maintained fixation until the end of the trial.

### Passive fixation task

Reference data for eye tracking quality from head-fixed animals were obtained from a passive fixation task. Animals were required to fixate a central fixation point for 100-140 ms. After this period, a grating stimulus was presented for 500 ms, and the animal was required to maintain fixation throughout the trial in order to obtain reward. Calibration was performed as in the head-free experiments and was kept identical across sessions.

### Psychometric analysis

We calculated hit rates and mean reaction times for all spatial frequency conditions and fitted the following four-parameter logistic model to the data (Cardillo, 2012):

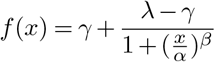

Where *γ* is the lower asymptote of the logistic function (fixed at chance performance of 12.5%, for the hit rate data, and restricted to be between 50-500ms for the RT data), and *λ* is the upper asymptote (restricted to be between 12.5-100% for the hit rate data, and between 50-500ms for the RT data). The *α* parameter corresponds to the inflection point and gives the spatial frequency value at which the hit rate, or mean RT, is halfway between the lower and upper asymptote. This parameter is also called the “acuity threshold”. The *β* parameter determines the steepness of the curve. Confidence intervals for RTs were calculated by bootstrapping (bias corrected and accelerated percentile method with 10,000 bootstrap replications). Confidence intervals for hit rates were calculated with the Clopper-Pearson method (Clopper and Pearson, 1934).

### Surgical procedures

A detailed account of all surgical procedures and recording methods will be described in a separate publication. In brief, animals were first implanted with a custom machined titanium head-post and a 3D printed titanium chamber. In a second surgery, silicon probes were semi-chronically implanted in areas V1 and V6 with one microdrive per area (Nano-Drive CN-01 V1, Cambridge NeuroTech, UK). Two 32-channel shanks with 250 µm spacing were implanted in V1, and four 32-channel shanks in V6 (H2 probe, Cambridge NeuroTech, UK). For experiments not described here, a viral vector (AAV1.CamKIIa.Chronos-eYFP-WPRE) was injected into area V6 just before electrode implantation. For monkey A, the stereotaxic coordinates from Paxinos et al. (2012) served as the anatomical guide for electrode implantation: The target coordinates for V1 were 8.5 mm caudal from the interaural line and 1.3 mm lateral from the midline, and the coordinates for V6 were 2.5 mm caudal from the interaural line and 3 mm lateral from the midline. For monkey D, we used a combination of Paxinos et al. (2012) and a CT-scan of the animal’s skull and chamber, to which an MRI template brain (Liu et al., 2018) was manually aligned: The target coordinates for V1 were 7.7 mm caudal from the interaural line and 1.3 mm lateral from the midline, and the coordinates for V6 were 2.6 mm caudal from the interaural line and 4.1 mm lateral from the midline.

Anesthesia was induced with an intramuscular (i.m.) injection of a mixture of alfaxalone (8.75 mg/kg) and diazepam (0.625 mg/kg). Tramadol (1.5 mg/kg) and metamizol (80 mg/kg) were injected i.m. for initial analgesic coverage. Subsequently, a continuous intravenous (i.v.) infusion was provided through the lateral tail vein to the animal. The i.v. mixture contained glucose, amino acids (Aminomix 1 Novum, Fresenius Kabi, Germany), dexamethasone (0.2-0.4 mg·kg^-1^·h^-1^), tramadol (0.5-1.0 mg·kg^-1^·h^-1^) and metamizol (20-40 mg·kg^-1^·h^-1^). The maximal infusion rate was 5 ml·kg^-1^·h^-1^. After ensuring appropriate depth of anesthesia, animals were placed in a stereotaxic frame. Animals were breathing spontaneously throughout the surgery via a custom face mask that applied isoflurane (0.5-2% in 100% oxygen). Heart rate, respiration rate and body temperature were constantly monitored (Model 1030 Monitoring Gating System, SAII, USA).

### Acquisition and processing of neuronal data

Neuronal signals were recorded through active, unity gain head stages (ZC32, Tucker Davis Technologies, USA), digitized at 24,414.0625 Hz (PZ2 preamplifier, Tucker Davis Technologies, USA) and re-sampled offline to 25 kHz. Sample-by-sample re-referencing was applied by calculating the median across all channels for each shank and subtracting this signal from each channel of the corresponding shank (Jun et al., 2017). Data was band-pass filtered with a 4th-order Butterworth filter (0.3-6 kHz) for spiking activity (as shown in Fig. 7a). For further analysis, multi-unit activity (MUA) was calculated by full-wave rectification, filtering with a 6th order low-pass Chebyshev-II filter (stopband attenuation of 50 dB) and down sampling to 1 kHz.

### Receptive field mapping and SNR analysis

All data for the receptive field mapping experiments were recorded under head-fixation. Data was first cut into epochs of 280 ms (from 100 ms before to 180 ms after stimulus onset) based on the onset timing of stimulus presentation as determined from the photodiode signal. For incomplete trials (break fixation trials), a given epoch was included in the analysis as long as the eye position remained inside the fixation window throughout the epoch. To reject artifacts, we calculated the standard deviation of MUA across time within each 280 ms epoch. Epochs in which the standard deviation was more than 10-times larger than the median standard deviation across all epochs were excluded from the analysis.

Recording sites were included in the analysis if the mean MUA from at least three different wedge stimuli and at least three different annulus stimuli evoked responses that were significantly larger than the MUA during baseline (paired t-test, alpha = 0.01). The baseline was defined as the MUA 100 ms to 0 ms before stimulus onset.

For the calculation of RFs, MUA from 0 ms to 100 ms after stimulus onset from all artifact-free epochs was averaged, and this value was multiplied with the 2D matrix containing the intensity values from the images shown at the corresponding epoch, separately for annuli and wedge stimuli. All the resulting images where summed up and divided by a bias image to normalize for unequal repetitions of images. The bias image was computed by summing up all images that were presented in all artifact-free epochs, separately for annuli and wedge stimuli. The final RF map was calculated by pixel-wise multiplication of the two maps and then scaled to range between zero and one.

Estimates of RF size and position were obtained by fitting a Gaussian function to the annulus data and a von-Mises function (the circular approximation of a Gaussian function) to the wedge data. Mean, and circular mean from the resulting model fits were used as peak eccentricity and polar angle, respectively. The RF size along the eccentricity axis (RFS_e_) was defined as the full width at half maximum (FWHM) of the Gaussian model fit. The RF size along the axis perpendicular to the eccentricity was calculated on the basis of the FWHM of the von-Mises model fit: The resulting circular arc length was used to calculate the corresponding chord length (RFS_c_). The RF size was defined as the geometric mean of RFS_e_ and RFS_c_.

For the analysis of signal-to-noise ratio (SNR), RF maps were computed as mentioned above, but repeatedly from a subset of the data. For each subsampling run, we randomly picked a small number of stimulus presentations per condition from the complete dataset. This procedure was performed 100 times for each repetition size step from one to nine repetitions. Nine repetitions were chosen as the maximum for the analysis because it was the lowest number of available repetitions in the dataset. The analysis was performed by using the neuronal data after stimulus presentation (0 to 100 ms) and, as a baseline control, by using data before stimulus presentation (−100 to 0 ms). The SNR was calculated as the amplitude ratio of signal and noise in decibels (dB). The signal was defined to be the mean value inside a region of interest (ROI) on the RF map. The noise was defined as the mean value outside the same ROI. The ROI was determined per recording site by thresholding the RF map that was calculated from all available data. Pixels with values larger than the half maximum of the RF map were defined as being within the ROI. For each recording site, the same ROI was used for all subsampling runs. Confidence intervals were calculated by bootstrapping (10,000 bootstrap replications) on the subsampled data. The mean SNR values and confidence intervals were averaged across animals and areas.

## Acknowledgements

PF acknowledges grant support by DFG (SPP 1665 FR2557/1-1, FOR 1847 FR2557/2-1, FR2557/5-1-CORNET, FR2557/6-1-NeuroTMR, FR2557/7-1 Dual-Streams), EU (HEALTH-F2-2008-200728-BrainSynch, FP7-604102-HBP, FP7-600730-Magnetrodes), a European Young Investigator Award, NIH (1U54MH091657-WU-Minn-Consortium-HCP), and LOEWE (NeFF). GR is supported by FAPESP (2017/10429-5 and 2018/16635-9). We thank Marianne Hartmann at the ESI for her continuous support in training the animals.

## Author contributions

PJ, FJK and PF designed research; PJ and FJK performed research; PJ and GR analyzed data; PJ, PF and GR wrote the paper. All authors edited and approved the final manuscript.

---

**Table 1:** Statistical tests reported in the Results. Values in the text are labeled by the letters in the left-hand column.

|  | Distribution | Type of test | Statistic | p value | Power | Sample size | Animals |
| --- | --- | --- | --- | --- | --- | --- | --- |
| a | Non-normal | Pearson correlation | $\rho = 0.145$ | $p = 0.0734$ | 0.435 | $n = 153$ | $n = 4$ |
| b | Normal | t-test (paired, two-sided) | $T = -19.803$ | $p = 0.00028$ | $\approx 1$ | $n = 4$ | $n = 4$ |
| c | Normal | t-test (paired, two-sided) | $T = 5.427$ | $p = 0.012$ | 0.935 | $n = 4$ | $n = 4$ |

**Extended Data Movie 1:**
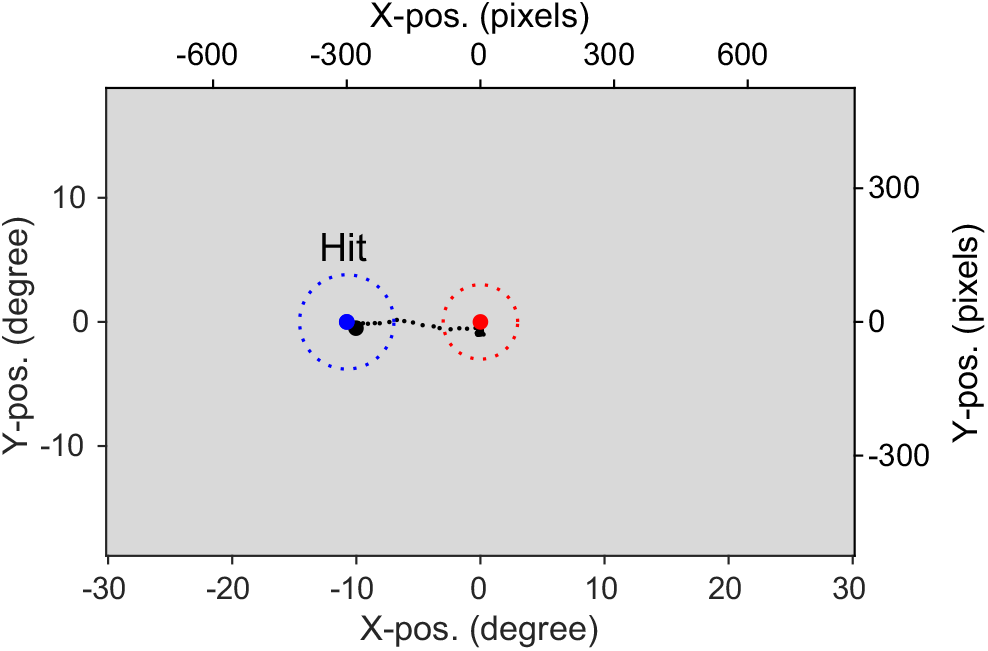
Head-free eye tracking during visual acuity measurements. Eye position and trial outcome from six example trials are shown in the movie (three hits and three miss trials). Red and blue dots indicate position of fixation point and target stimulus, respectively. Dashed circles are the corresponding eye tracking windows. Small black dots show the last 100 ms of eye position during the trials. Large black dot shows current eye position.

**Extended Data Figure 3-1:**
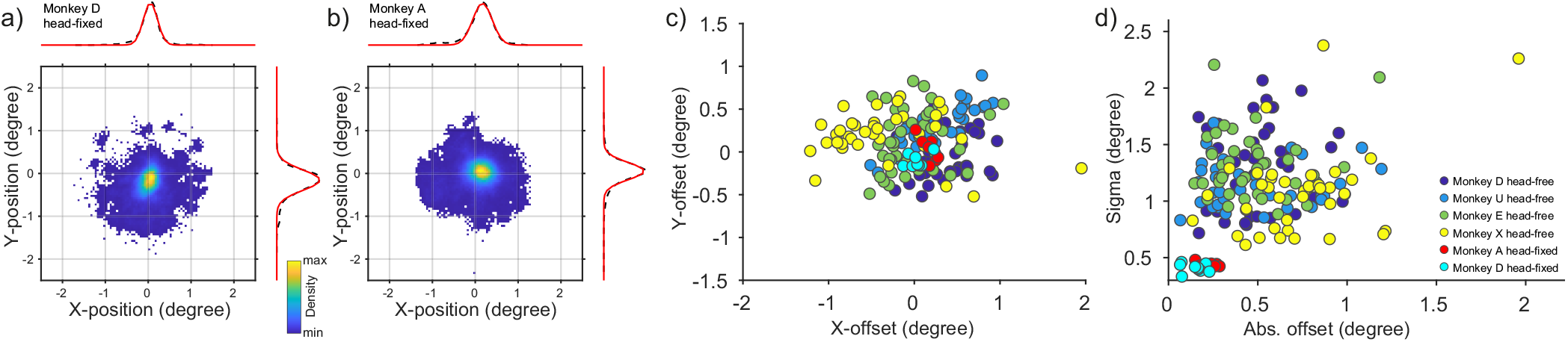
Quality of eye signals, head-free versus head-fixed. **a**,**b)** Example density plots of eye position during passive fixation for monkeys A and D. Data are taken from time of fixation onset until the end of the trial. Dashed lines show the average density for X-and Y-position. Red lines show Gaussian fits. **c)** X-and Y-offset values for all head-free (n = 153) and head-fixed (n = 17) sessions, color coded as indicated in panel d). **d)** Scatter plot of offset versus sigma values for head-free and head-fixed sessions.

**Extended Data Figure 7-1:**
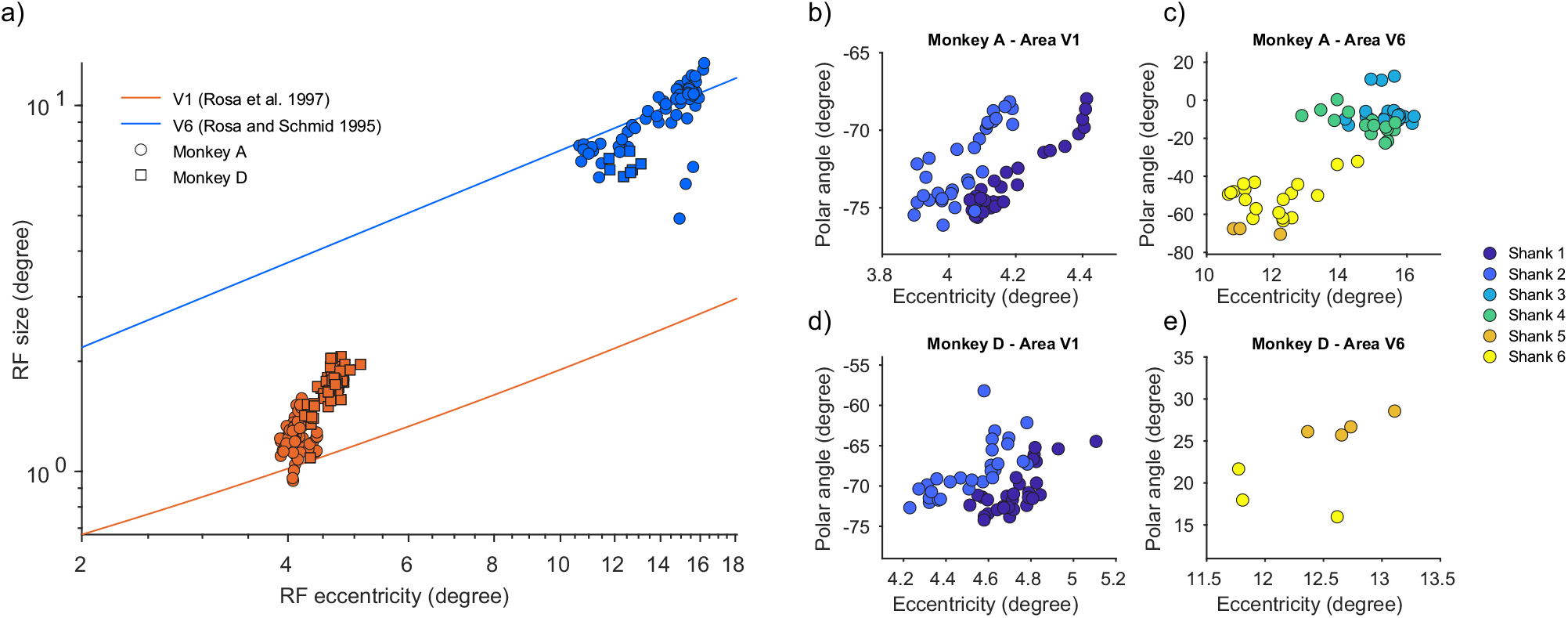
Receptive field sizes and positions. **a)** Receptive field size as a function of eccentricity from all stimulus-driven recording sites in areas V1 and V6. **b-e)** Receptive field positions from all stimulus-driven recording sites, separately for each monkey and area. Color indicates on which electrode shank the recording site was located. Note the clear clustering of positions within the shanks. Because animals were still participating in experiments at the time of this this work, no histological data on the electrode shank placement was available. Thus, the progression of RF positions across shanks should be interpreted with care.

